# Evoregions: Mapping Shifts in Phylogenetic Turnover Across Biogeographic Regions

**DOI:** 10.1101/650713

**Authors:** Renan Maestri, Leandro Duarte

**Author notes:** Correspondence be sent to: Departamento de Ecologia, Universidade Federal do Rio Grande do Sul, Av. Bento Gonçalves 9500, CP 15007, Porto Alegre RS 91501-970, Brazil.

## Abstract

Biogeographic regionalization offers context to the geographical evolution of clades. The positions of bioregions inform both the spatial location of clusters in species distribution and where their most important boundaries are. Nevertheless, defining bioregions based on species distribution alone only incidentally recovers regions that are important during the evolution of the focal group. The extent to which bioregions correspond to centers of independent diversification depends on how clusters of species composition naturally reflect the radiation of single clades, which is not the case when mixed colonization occurred. Here, we showed that using phylogenetic turnover based on fuzzy sets, instead of species composition, led to more adequate detection of evolutionary important bioregions, that is, regions that truly account for the independent diversification of lineages. Mapping those evoregions in the phylogenetic tree quickly reveals the timing and location of major shifts of biogeographic regions. Moreover, evolutionary transition zones are easily mapped, and permits the recognition of regions with high phylogenetic overlap. Our results using the global radiation of rats and mice (Muroidea) recovered four evoregions—three major evolutionary arenas corresponding to the Neotropics, a Nearctic-Siberian, and a Paleotropical-Australian evoregion, and a fourth and fuzzy Afro-Palearctic evoregion. In comparison, an analysis with a method considering species distribution alone found 52 bioregions. Evoregions is a useful framework whenever the question is related to the identification of the most important centers of a group’s diversification history and its evolutionary transitions zones.

## Introduction

Biogeographic boundaries reflect important geographic limits during the evolutionary history of clades (Wallace 1876). Boundaries divide the world into regions of endemicity, since lineages of a focal clade are thought to have evolved in isolation from other such lineages into each region. These biogeographic discrete units compose the so-called biogeographical realms, regions, dominions, or provinces (Holt et al. 2013; Morrone 2014; Costello et al. 2017), named depending on the spatial scale of the study (see Morrone 2015 and Vilhena and Antonelli 2015 for a discussion about terminology). Each monophyletic clade (taxa) is likely to have its unique set of important biogeographic regions (bioregions), reflecting the principal geological and climatological factors in action during the timing of diversification, and the particular organism’s dispersal abilities (Edler et al. 2017; Maestri et al. 2019). Boundaries for different taxa that are found later to be in coincident position may help to infer global bioregions. However, knowing the evolutionary important bioregions for single monophyletic clades may be more informative, and less artificial, then trying to resume very different histories together.

Bioregions can be defined in various ways, from using expert knowledge to more recent data-driven approaches (Kreft and Jetz 2010; Holt et al. 2013; Olivero et al. 2013; Vilhena and Antonelli 2015; Edler et al. 2017). Frequently, data-driven methods gather a matrix of presence/absence of species across assemblages, usually cells in a grid, and apply a quantitative procedure—as species turnover, network, or cluster analysis — to assembly cells into bioregions. In common, virtually all approaches (i) use the dissimilarity in species composition alone, without considering phylogenetic relationships among taxa, and (ii) seldom account for biogeographic transition zones. Bioregions demarcated using species distribution may find regions of endemicity defined by multiple colonization of species belonging to various phylogenetic lineages, and therefore lack single histories of diversification. Holt et al. (2013) made a first attempt to classify global bioregions based on phylobetadiversity patterns. Their approach used the Simpson index of beta-diversity to quantify the sharing of tree branches among assemblages (Holt et al. 2013). However, such simplified approach relying on counting of branches may not be so informative when multiple clades with different histories are grouped together, causing the identification of bioregions attributed to single diversification events where in fact those regions resulted from multiple colonization/diversification events (Kreft and Jetz 2013). Furthermore, spatial scale and geographic distances can artificially influence beta-diversity metrics of turnover (Vellend 2001; Vilhena and Antonelli 2015), and such metrics also do not fully account for phylogenetic distances and phylogenetic imbalance (Leibold et al. 2010; Kreft and Jetz 2013; Duarte et al. 2016). To identify and account for transition zones, a promising approach using fuzzy logic has been proposed by Olivero et al. (2013), which captures better the intricacies of species distribution patterns, but such approach has never been extended to incorporate phylogenetic relationships among taxa. For all these reasons, the development of suitable approaches to delimit bioregions remains an open avenue in historical biogeography.

The identification of biogeographic regions that consider the differences in evolutionary history among species continues to be a challenge to biogeographers. Post-hoc approaches based on ancestral range estimation have been used to find evolutionary relationships among bioregions defined as biogeographic units sharing common species distribution patterns (Ree and Smith 2008), and/or seek for the historical and ecological drivers of bioregion boundaries (Ficetola et al. 2017). In the end, what biogeographic regions really need to represent are the histories of independent diversifications that occurred within the region, and this is only indirectly accomplished using species composition. An approach that simultaneously considers evolutionary distances among taxa and among assemblages might extend the definition of bioregion in order to incorporate evolutionary relationships among taxa.

In this study, we introduce the concept of evoregion as a biogeographic region where most of the resident species stem from one or a few *in situ* radiations. Further, biogeographic regions showing high phylogenetic turnover, and therefore having a low affiliation to a single evoregion, can be defined as evolutionary transition zones. We propose a fuzzy logic-based approach to classify evoregions and their respective evolutionary transition zones. Our approach considers both pairwise phylogenetic divergences among taxa and tree imbalance (Pillar and Duarte 2010; Duarte et al. 2016), and therefore permits a complete assessment of evolutionary divergences between biogeographic regions. The evoregion approach allows (i) to map the geographic regions where the main diversification events for a given clade occurred, (ii) to characterize evolutionary transition zones, that is, biogeographic regions showing high phylogenetic turnover, and (iii) to assess the timing and approximate position of major evoregion shifts in the phylogenetic tree.

We illustrate the application of the evoregion concept by evaluating the worldwide evoregions of Muroidea, which is a group of rats and mice with over than 1,600 species distributed throughout the globe, and represents more than a quarter of all mammalian diversity (Wilson and Reeder 2005; Burgin et al. 2018). The group likely originated in Eurasia (Qiu and Li 2003; Jansa et al. 2009), and its richest subclades emerged during the independent radiations of Cricetidae—and its principal subfamily Sigmodontine—and Muridae, starting in the Miocene (Musser and Carleton 2005; Steppan and Schenk 2017; Burgin et al. 2018). Knowledge of the most important centers for muroids diversification remains an open question. Historically, Muridae occupation is associated with an extensive Old-World region including Eurasia and the Paleotropics and adjacent islands, while the Cricetidae non-sigmodontines (Cricetinae, Arvicolinae, Neotominae, and Tylomyinae) are widespread across the Holarctic region, and Sigmodontinae diversified in the Neotropics (Steppan et al. 2004; Jansa et al. 2009; Fabre et al. 2012). From their diversification history as inferred from the phylogenies and paleontological records in association with scattered evidences of biogeographical distribution (Conroy and Cook 1999; Jansa and Weksler 2004; Musser and Carleton 2005; Fabre et al. 2012; Schenk et al. 2013), at least these three major biogeographic regions of diversification can be expected as the main centers of independent diversification: (i) Paleotropical region including Africa, Southeast Asia and Australasia, (ii) Holarctic region including Eurasia and North America, and (iii) the Neotropics. Other events of independent divergence include Nesomyidae in Africa and its Malagasy subfamily Nesomyinae, and Australasia radiations of Murinae subclades. How many of those or other regions would be recovered as important bioregions for the study group using a data-driven approach? Here, we compared evoregions with bioregions defined using only species composition (Edler et al. 2017). We predict that evoregions will find sharpest bioregions, fewer in number but clearly related to *in situ* diversification events, than a method based on species composition.

## Material and Methods

### Muroidea Data

Species distributions for 1473 living species of Muroidea were taken from the IUCN database (IUCN 2008). Presence/absence of species was calculated over a global grid map with 4,161 cells of 2°x 2° degrees of latitude and longitude, which is an appropriate scale for global studies (Hurlbert and Jetz 2007). Presence on each cell was assigned if at least 25% of the cell was covered by a species range. A phylogenetic tree for Muroidea was taken from Steppan and Schenk (2017), the most comprehensive and up to date phylogeny for the group (the outgroup Dipodoidea was excluded from the tree). After pruning both datasets for coincident species, 670 species with both incidence data and phylogenetic information were retained for further analyses, which corresponds to more than 40% of taxonomic species diversity and nearly 75% of species with current phylogenetic information available.

### Phylogenetic Turnover

We define phylogenetic turnover as variation in the phylogenetic composition of Muroidea among grid cells, which was measured with the phylogenetic fuzzy-weighting method (Pillar and Duarte 2010). This method accounts for phylogenetic distances and tree imbalance, and shows greater statistical performance — higher power and increased effect sizes in statistical analyses — than other metrics of phylogenetic beta-diversity (Duarte et al. 2014, 2016). Accordingly, pairwise phylogenetic covariances between murid species were standardized within columns, resulting in a matrix **Q** depicting degrees of phylogenetic belonging of each species to every other species (Duarte et al. 2016). The pairwise degree of phylogenetic belonging of species *i* to species *j* captures, in a single value, the amount of phylogenetic divergence between *i* and *j*, and also the rate of diversification between *i* and the ancestral node δ connecting *i* to *j*. If the path linking species *i* to δ shows a higher rate of diversification than the path connecting *j* to δ, then species *i* will show a lower degree of belonging to *j* than *j* to *i* (Duarte et al. 2014, 2016). Therefore, matrix **Q** is a phylogenetic dissimilarity matrix that expresses, simultaneously, symmetric phylogenetic divergences among species and also asymmetric diversification trajectories connecting them. The method is described in detail in Duarte et al. (2016). Matrix **Q** was then multiplied by a matrix that describes species occurrences (presence/absence) in the cells, resulting in matrix **P**, which describes phylogeny-weighted species composition (or simply phylogenetic composition) for a set of assemblages. Differences in phylogenetic composition among cells express phylogenetic turnover. Both **P** matrix and PCPS were computed in the R statistical environment (R Core Team 2018) using the package *PCPS* (Debastiani and Duarte 2014).

### Evoregions

To map phylogenetic turnover across the cells we used Discriminant Analysis of Principal Components based on k-means non-hierarchical clustering (DAPC; Jombart et al. 2010). DAPC is based on principal components extracted from the input data; nonetheless, phylogenetic turnover among cells are hardly linear, which is an assumption of Principal Component Analysis (Pielou 1984). Empirical evidence has shown that non-Euclidean resemblance measures, such as Bray-Curtis dissimilarities are well suited for analyses of ecological data (Legendre and Anderson 1999; Legendre and Legendre 2012). Thus, prior to DAPC we computed Principal Coordinates of Phylogenetic Structure (PCPS; Duarte 2011) from matrix **P**, which implies performing Principal Coordinate Analyses on matrix **P** based on square-rooted Bray-Curtis dissimilarities between cells. PCPS eigenvectors capture gradients of phylogenetic turnover across grid cells (Duarte et al. 2016). DAPC was then performed taking all PCPS eigenvectors as input data, but using only those principal components containing ≥ 5% of total information of **P** for discriminant analysis. This procedure was based on the same basic principle of distance-based Redundancy Analysis (db-RDA; Legendre and Anderson 1999). The optimal number of groups obtained from such distance-based DAPC was defined finding the sharpest decrease in successive Bayesian Information Criterion (BIC) values (Jombart et al. 2010), computed for a set of group sizes ranging from two to 12 groups. We followed the procedure proposed by (Jombart et al. 2010) to determine the optimal group size involved 1) performing Ward’s clustering of BIC values; 2) splitting BIC values into two groups; 3) finding the highest BIC value among the group containing the lowest BIC values. The group size corresponding to such BIC value was considered as the optimal number of evoregions for Muroidea. DAPC was performed using the function ‘find.clusters’ of the R package *adegenet* (Jombart 2008).

### Evolutionary Transition Zones

As we have seen above, evoregions are defined based on a clustering algorithm that allocates each cell to a given group. Nonetheless, the degree of membership of each cell to its respective evoregion may vary within a given evoregion. Mapping the degree of membership of cells to its evoregion is a manner to visualize biogeographic transition zones (Olivero et al. 2013). The degree of membership of a cell to its respective evoregion was computed as the mean phylogenetic dissimilarity of the cell to all other cells belonging to the same evoregion (Olivero et al. 2013), rescaled to vary between 0 and 1. Values closer to unity indicate a higher degree of membership (affiliation) of a given cell to its respective evoregion.

### Reconstructing Ancestral Affiliation of Clades to Evoregions

A major advantage of phylogenetic fuzzy weighting when compared to other methods for evaluating phylogenetic turnover relies on the possibility of assessing the affiliation of each species not only to its ancestral clades, but also to evoregions where they occur. The affiliation of species to evoregions allows reconstructing shifts in the distribution of lineages across evoregions, which make possible inferring evolutionary shifts of clades across the biogeographic space.

Of course, a single species can occur simultaneously in two or more evoregions, which imposes a problem for reconstructing the affiliation of ancestral clades to evoregions. One way to resolve this issue is to use explicit biogeographic models that allow taxa to occur in two or more areas simultaneously (Ree and Smith 2008). This explicit biogeographic approach is recommended, and is particularly important when the main objective is to locate the ancestral ranges. Here, since we are more interested in observe single terminal colors (i.e. connect a tip to a unique evoregion) in order to visualize how many members of a single clade are present in an evoregion, we used a simpler approach. To overcome this issue, we allocated each species to a single evoregion, which was accomplished considering the evoregion where a given species most frequently occurs. For instance, if 70% of the cells where species *i* occurs are allocated to evoregion 1, and 30% in evoregion 2, we can assume there is a probability *p* = 0.7 of species *i* to belong to evoregion 1, and *p* = 0.3 of belonging to evoregion 2. If we assume 0.7 as a reasonable threshold to consider a species as belonging to a given evoregion, whenever the frequency of occurrence of a species to a given evoregion reaches this threshold, we allocate it to evoregion 1. Otherwise, if that species never reaches the threshold frequency, it cannot be allocated to a single evoregion. In that case, it should be classified as a widespread species, which is a separate category. We tested thresholds varying between 0.6 and 0.9, which showed similar results. Here we present only the results for threshold = 0.7, but the decision on the threshold value is arbitrary.

After defining the affiliation of all species to a single evoregion (or classifying it as widespread), we proceeded to the estimation of ancestral evoregions along the Muroidea phylogenetic tree using stochastic character mapping based on an equal-rates model for the transition matrix of our discrete traits (1000 simulations), which is implemented in the function ‘make.simmap’ of the R package *phytools* (Revell 2012). The 1000 character maps allowed us to estimate and plot uncertainty on history estimation.

### Comparison with Bioregions Defined Using Species Composition

A novel and promising method to identify key bioregions based on data on species composition, instead of on expert opinion, is the Infomap Bioregions routine (Edler et al. 2017). Briefly, this approach uses a clustering algorithm on a bipartite network of species occurrences within cells to classify bioregions. We imputed all the range maps for Muroidea into the Infomap website (https://bioregions.mapequation.org/), and generated bioregions using a cell size of 2°x 2° degrees of latitude and longitude to allow direct comparison to evoregions. The minimum cell capacity was set to 1 species. The resulting shapefile containing the bioregions is made available in the Supplementary Material.

## Results

Distance-based DAPC indicated the occurrence of four distinct evoregions for Muroidea across the globe (fig. 1). In the Northern Hemisphere, we observe the occurrence of two main evoregions: Nearctic-Siberian and Afro-Palearctic, with the first characterizing mainly the Nearctic region in the Western hemisphere and parts of the Siberia in the Eastern hemisphere, while the latter comprises the Palearctic region extending from Europe to East Asia and including northern parts of Africa and the Middle East, but also containing southern parts of Africa and Madagascar. Across the Southern Hemisphere, two evoregions are dominant: Paleotropical-Australian and Neotropical, where the first comprises tropical parts of Africa, Southeastern Asia and Australasia, and the latter is clearly Neotropical.

**Figure 1:**
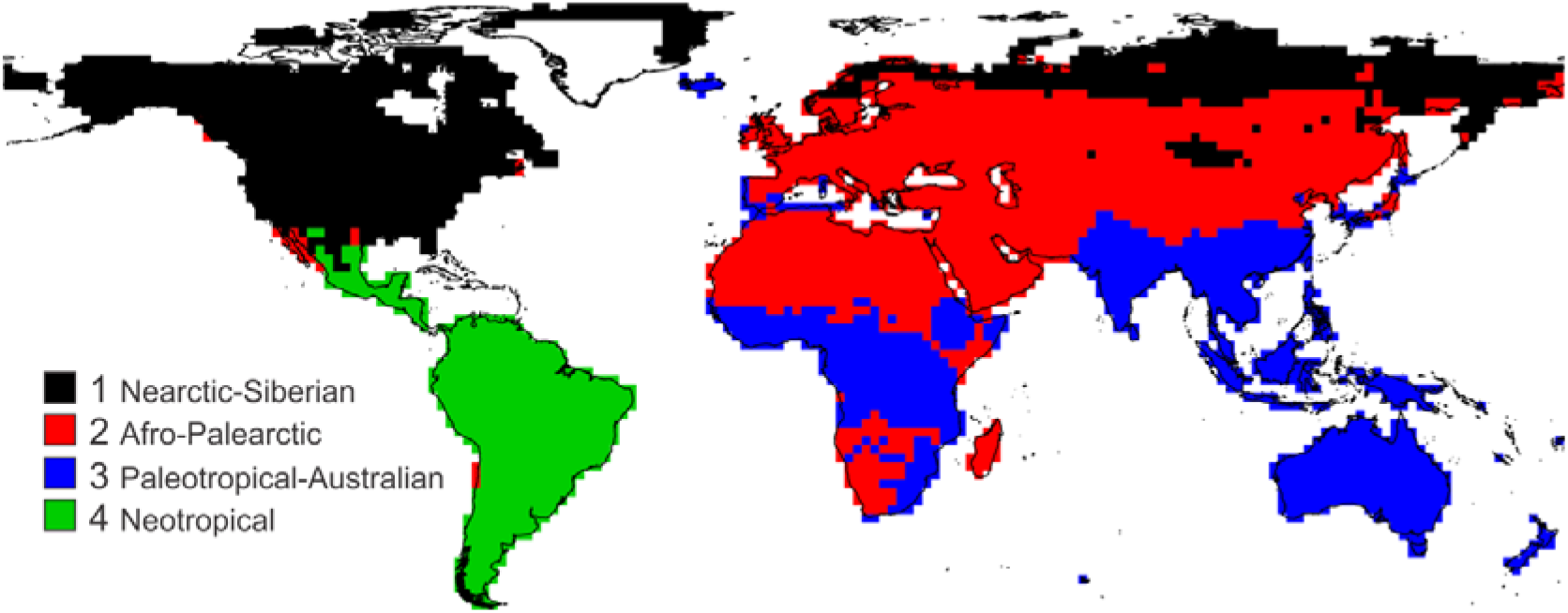
Evolutionary important biogeographic regions—evoregions—for Muroidea. Colors denote different evoregions. Evoregions were constructed based on phylogenetic turnover, see the main text for further information.

In figure 2 we can observe the occurrence of three large biogeographic transition zones for Muroidea clades: one in Central America and southern parts of North America in the boundary between the Nearctic-Siberian and the Neotropical evoregion; a second covering African parts of the Afro-Palearctic evoregion (northern Africa and the Middle East, and southern Africa and Madagascar) and parts of Asia in the boundaries with the Nearctic-Siberian region; and a third in southeastern China within the Paleotropical-Australian evoregion. On the other hand, most assemblages of the Neotropical evoregion and of the Paleotropical-Australian evoregion showed high values of affiliation, indicating high phylogenetic homogeneity within those evoregions.

**Figure 2:**
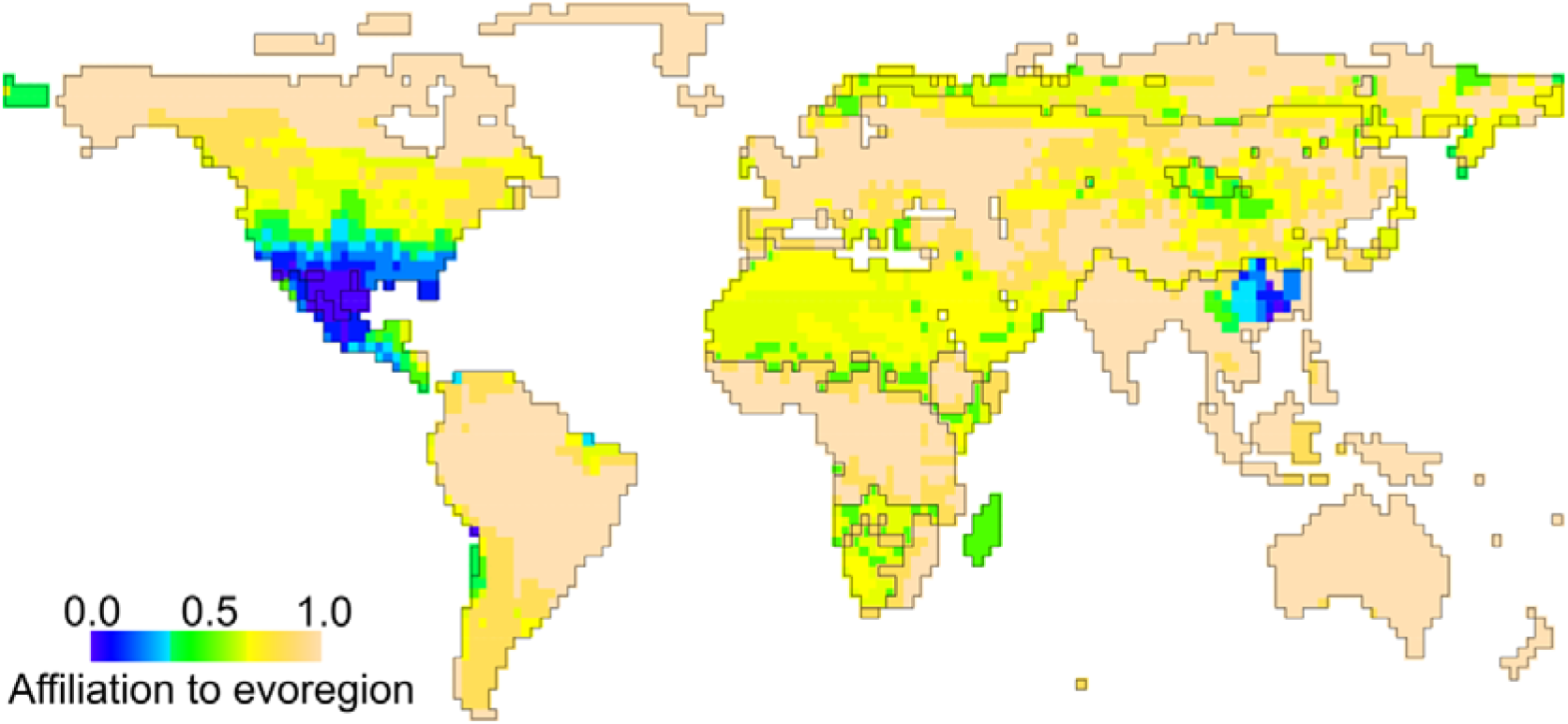
Evolutionary transition zones among evoregions. The map informs how affiliated each cell is to its evoregion—smaller values indicate low affiliation and therefore high phylogenetic turnover typical of transition zones. A shapefile of the evoregions was overlaid on the map: the contours depict the boundaries among evoregions (see fig.1).

The Neotropical evoregion is determined by the *in situ* diversification of sigmodontine rodents (Muroidea, Cricetidae, Sigmodontinae), making it the evoregion most clearly defined by a single radiation (fig. 3). The other major subfamilies in Cricetidae: Neotominae (Nearctic) and Arvicolinae (mostly Palearctic), occur through the wide Holarctic region, and lineages of both subfamilies diversified over (and define) the Nearctic-Siberian evoregion. Lineages of Arvicolinae and other cricetids, as well as lineages of Muridae and of some clades stemming from the most basal nodes in Muroidea phylogeny (including Nesomyinae rodents from Madagascar) grouped in the wide Afro-Palearctic evoregion (fig. 3). However, large groups of assemblages within this latter evoregion have low affinity to it, due to high phylogenetic turnover (fig. 2). The Paleotropical-Australian evoregion is in large part determined by the *in situ* diversification of Muridae lineages (fig. 3). Note that the largest part of the species are classified into an evoregion along with its close relatives (fig. 3), exemplifying the concept of evoregion as a biogeographic region where most of the resident species stem from one or a few *in situ* radiations.

**Figure 3:**
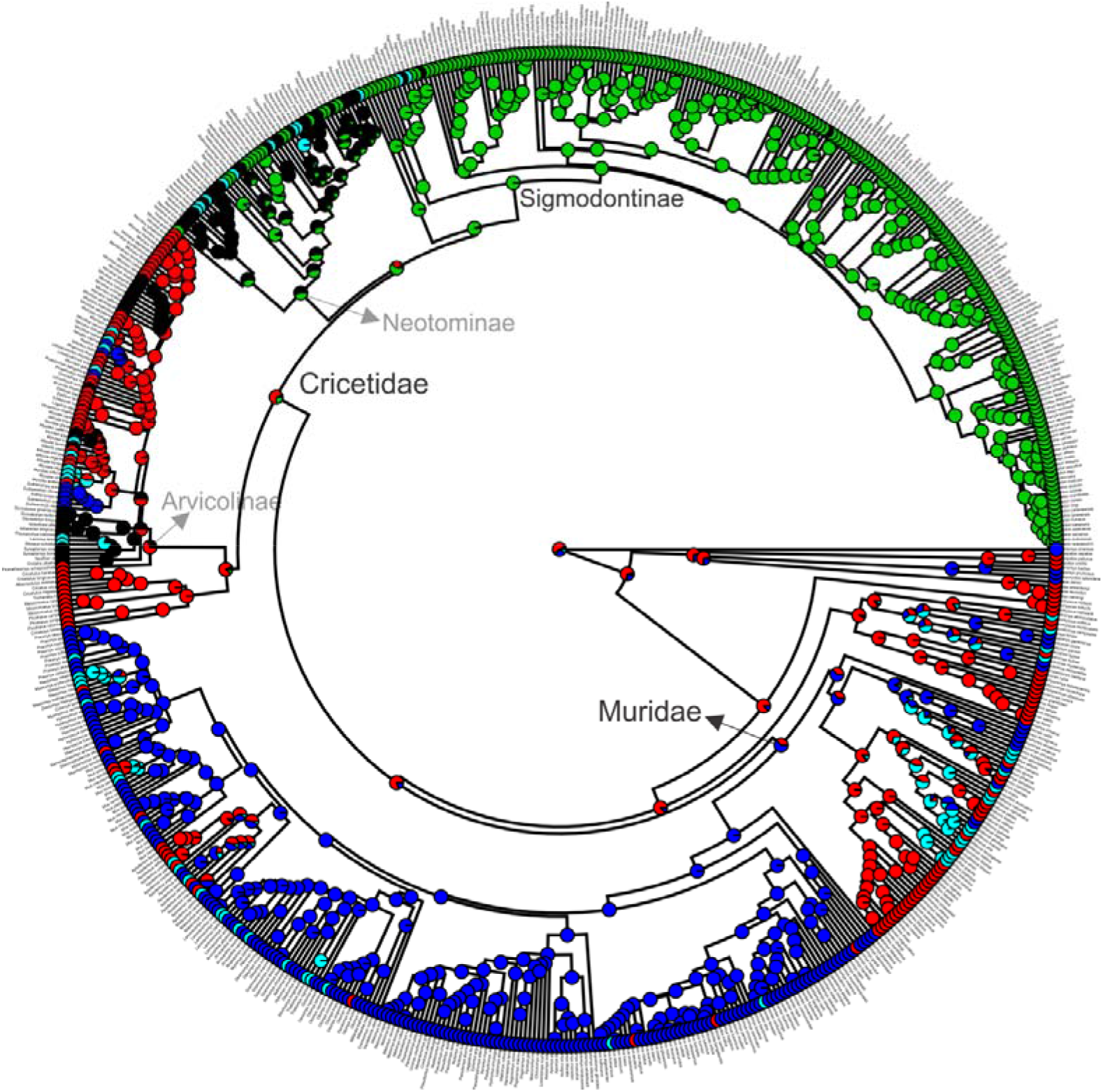
A phylogenetic tree of Muroidea with terminal colors representing the predominant evoregion for each species. Species were considered to belong to an evoregion if 70% or more of its distribution lied within a single evoregion (threshold = 0.7). Widespread species (those below the threshold) appear in cyan. Colors are the same as in fig. 1. Ancestral evoregions are a summary of 1000 stochastic character histories estimated using an equal-rates transition matrix.

The Infomap bioregion method identified 52 bioregions based on species composition (fig. 4). Some of the Infomap bioregions have a clear correspondence to an evoregion, for example, with the North American evoregion, northern parts of the Afro-Palearctic evoregion, and parts of the Paleotropical-Autralian evoregion (compare North Amercia, the north of Africa, the sub-Sahara Africa, and Australia in figs. 1 and 4). Overall, it is noticeable that many boundaries identified with evoregions are also present in the Infomap bioregions. The main difference between both approaches is that Infomap bioregions splitted the world in more and smaller parts. For example, all Neotropical region was identified as a single evoregion (to the exception of a few cells in west Chile), while seven bioregions were identified in the Neotropics with Infomap. Nevertheless, the boundary between the Neotropics and North America was identified in the approximate same position with both approaches.

**Figure 4:**
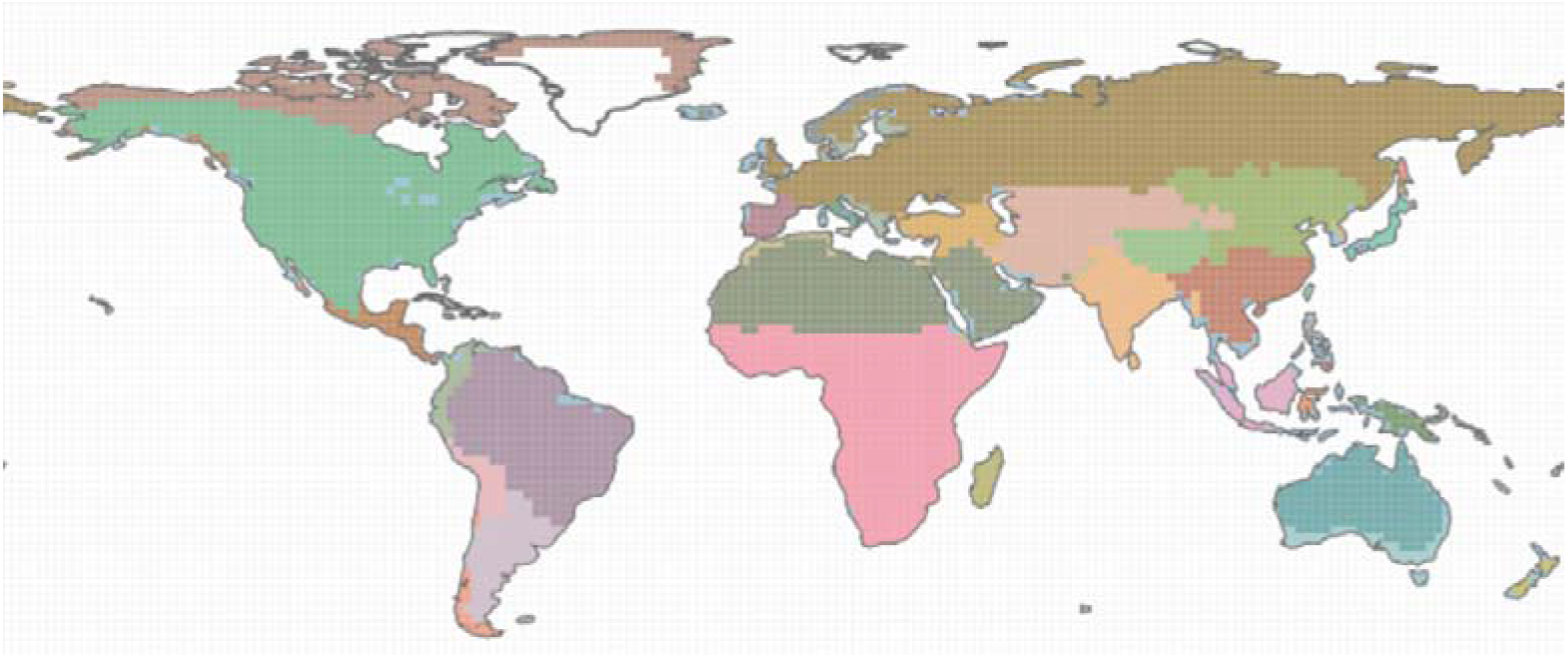
Infomap bioregions for Muroidea. Range maps were used to find the bioregions considering a cell size of 2°×2°. Colors denote the 52 different bioregions identified.

## Discussion

The radiation of a single monophyletic clade that generates high endemism in a region is the core concept in a search for evolutionarily important — or phylogenetically distinct — bioregions (Holt et al. 2013; Kreft and Jetz 2013). Of course, species distributions are messy and do not follow a simple one-to-one relationship between clade and geographic region, which calls for an approach that is able to classify evolutionary histories as best as possible into bioregions. Our classification of assemblages into evoregions achieves that goal for Muroidea since each evoregion can be interpreted as a coherent region where members of unique lineages diversified mostly *in situ*. This same rationale cannot be easily applied to bioregions defined using species composition, for example. Therefore, this is the first major advantage of using evoregions instead of bioregions based on species composition: evoregions capture the geographic history of independent diversification of lineages within the focal clade of interest, and the boundaries dividing bioregions are likely to be important for lineage splitting. Moreover, advancing over metrics of beta-diversity based on counting of tree branches (Holt et al. 2013; Kreft and Jetz 2013), by considering phylogenetic distances the evoregions adequately detect and classify young and rapid radiations that occurred *in situ* into unique evoregions (e.g., Sigmodontinae and Muridae lineages defining the Neotropical and Paleotropical-Australian evoregions, respectively), while a method using counting of branches would be unable to segregate adequately such rapid radiations from the older radiations that compose the northern (and fuzziest) evoregions.

Notwithstanding, macroevolutionary dynamics of speciation, extinction, and dispersal are complex, especially in continents (Albert et al. 2017). Evoregions serve as a useful simplification considering the limitations of any other arbitrarily constructed bioregion (Kreft and Jetz 2010) in terms of difficulty to assign all species or clades unequivocally to a single bioregion, but with the advantage of having a strong evolutionary basis. Events of regional colonization by members of multiple phylogenetic lineages are common in nature, and we should expect a blurring in the assignment of evoregions. This is when it comes to the second major advantage of using evoregions: the boundaries between evoregions can easily be perceived as transition zones, a necessary step to the understanding of limits among evoregions and their overall reliability. For muroids, mapping evolutionary transition zones highlight regions of high phylogenetic turnover and also identify the geographic areas with low reliability of belonging to an evoregion (e.g. parts of Africa and Madagascar in the Afro-Palearctic evoregion, fig. 2). Moreover, by comparing map information on transition zones with the reconstruction of evoregions on the tree (compare figs. 2 and 3) it is evident which lineages contribute the most to the high phylogenetic turnover in these regions. For example, the boundary between the Nearctic-Siberian and the Neotropical evoregions shows assemblages with low affinity in Central America, which is evident as well in the phylogenetic tree, where members of Neotominae may occupy one or the other evoregion.

The identified evoregions are areas of major diversification events for Muroidea, and at once provide the information that is usually gathered using independently defined bioregions together with ancestral area estimation and analysis of diversification. Our results for Muroidea are qualitatively similar to that of Schenk et al. (2013), for example, which used ancestral range estimation on seven areas defined *a priori* (the areas were defined by coupling information on tectonics history and conventional biological realms along with diversification analysis for each region). However, using an *a priori* definition of bioregions, as well as defining bioregions based on the data on species distribution alone, led to the assignment of more bioregions than would be necessary, assuming that evoregions, by definition, find only the sharpest regions of diversification. That could led to an overestimation of the number of biogeographic transition events: many transitions were detected among Southeastern Asia, Sahul, and parts of Africa using an expert definition of bioregions (Schenk et al. 2013), and presumably many more would be detected if 52 bioregions were considered. On the other hand, evoregions classified a single Paleotropical-Australian region as a unique diversification arena without boundaries within it. Considering a whole-tree scale, a single Paleotropical region (including Australasia) makes sense given the close phylogenetic relationship and recent diversification events of Murinae lineages that are widespread through this evoregion. Evoregions can then serve as the most natural diversification areas for further analysis of diversification dynamics.

In summary, while using expert-based defined bioregions or bioregions defined using species composition can lead to a more refined partition of bioregions (Kreft and Jetz 2010; Edler et al. 2017), it can also lead to the detection of many biogeographic transitions that are unreal from an evolutionary standpoint—biogeographical regions lacking a coherent history of diversification considering the phylogenetic scale under investigation. If needed, refined evoregions can be achieved by further applying the framework described here to each of the identified lineages with unique histories (fig. 3), leading to a hierarchical classification of evolutionary bioregions. This could bring evolutionary rigor into the hierarchical classification of bioregions, approximating it from the hierarchical taxonomic classification based on phylogenetics. In addition, by applying evoregions to multiple independent and world-wide distributed clades it would be possible to summarize the findings to achieve novel classifications of global biogeographic realms.

Certainly, biogeographic classifications based solely on species composition are of great value, especially for conservation purposes where species uniqueness alone is important, and/or whenever the aim is to find regions of endemicity regardless of the diversification history. Evoregions is an alternative for crucial questions related to historical biogeography and macroevolution, when knowing the geographical history of diversification is the ultimate goal. Moreover, the ability to identify evolutionary transition zones and areas of high and low affinity to an evoregion permits a better assessment of the intricate distribution of species, and aid to a careful interpretation of biogeographic regions.

## Acknowledgments

RM thanks to CAPES and CNPq (grant 406497/2018-4) for funding. LD research activities have been supported by CNPq Productivity Fellowship (grant 307527/2018-2). LD research has been conducted in the context of the National Institutes for Science and Technology (INCT) in Ecology, Evolution and Biodiversity Conservation, supported by MCTIC/CNPq (proc. 465610/2014-5) and FAPEG.

## Literature Cited

Albert, J. S., D. R. Schoolmaster, V. Tagliacollo, and S. M. Duke-Sylvester. 2017. Barrier displacement on a neutral landscape: Toward a theory of continental biogeography. Systematic Biology 66:167–182.

Burgin, C. J., J. P. Colella, P. L. Kahn, and N. S. Upham. 2018. How many species of mammals are there? Journal of Mammalogy 99:1–14.

Conroy, C. J., and J. A. Cook. 1999. MtDNA evidence for repeated pulses of speciation within arvicoline and murid rodents. Journal of Mammalian Evolution 6:221–245.

Costello, M. J., P. Tsai, P. S. Wong, A. K. L. Cheung, Z. Basher, and C. Chaudhary. 2017. Marine biogeographic realms and species endemicity. Nature Communications 8:1–9.

Debastiani, V. J., and L. D. S. Duarte. 2014. PCPS – an R-package for exploring phylogenetic eigenvectors across metacommunities. Frontiers of Biogeography 6:144–148.

Duarte, L. D. S. 2011. Phylogenetic habitat filtering influences forest nucleation in grasslands. Oikos 120:208–215.

Duarte, L. D. S., R. S. Bergamin, V. Marcilio-Silva, G. D. D. S. Seger, and M. C. M. Marques. 2014. Phylobetadiversity among forest types in the Brazilian Atlantic Forest complex. PLoS ONE 9:1–10.

Duarte, L. D. S. S., V. J. Debastiani, A. V. L. L. Freitas, and V. D. Pillar. 2016. Dissecting phylogenetic fuzzy weighting: theory and application in metacommunity phylogenetics. Methods in Ecology and Evolution 7:937–946.

Edler, D., T. Guedes, A. Zizka, M. Rosvall, and A. Antonelli. 2017. Infomap Bioregions□: Interactive Mapping of Biogeographical Regions from Species Distributions. Systematic Biology 66:197–204.

Fabre, P.-H., L. Hautier, D. Dimitrov, E. J. P. Douzery, P. Douzery, and J. Emmanuel. 2012. A glimpse on the pattern of rodent diversification: a phylogenetic approach. BMC evolutionary biology 12:88.

Ficetola, G. F., F. Mazel, and W. Thuiller. 2017. Global determinants of zoogeographical boundaries. Nature Ecology & Evolution 1:89.

Holt, B. G., J.-P. Lessard, M. K. Borregaard, S. A. Fritz, M. B. Araújo, D. Dimitrov, P.-H. Fabre, et al. 2013. An Update of Wallace’s Zoogeographic Regions of the World. Science 339:74–78.

Hurlbert, A. H., and W. Jetz. 2007. Species richness, hotspots, and the scale dependence of range maps in ecology and conservation. Proceedings of the National Academy of Sciences USA 104:13384–13389.

IUCN. 2008. IUCN Redlist of Threatened Species.

Jansa, S. A., T. C. Giarla, and B. K. Lim. 2009. the Phylogenetic Position of the Rodent Genus Typhlomys and the Geographic Origin of Muroidea. Journal of Mammalogy 90:1083–1094.

Jansa, S. A., and M. Weksler. 2004. Phylogeny of muroid rodents: Relationships within and among major lineages as determined by IRBP gene sequences. Molecular Phylogenetics and Evolution 31:256–276.

Jombart, T. 2008. adegenet: a R package for the multivariate analysis of genetic markers. Bioinformatics 24:1403–1405.

Jombart, T., S. Devillard, and F. Balloux. 2010. Discriminant analysis of principal components: a new method for the analysis of genetically structured populations. BMC Genetics 11:1–15.

Kreft, H., and W. Jetz. 2010. A framework for delineating biogeographical regions based on species distributions. Journal of Biogeography 37:2029–2053.

Kreft, H., and W. Jetz. 2013. Comment on “ An Update of Wallace’s. Science 341:9–10.

Legendre, P., and M. J. Anderson. 1999. Distance-based redundancy analysis: testing multispecies responses in multifactorial ecological experiments. Ecological Monographs 69:1–24.

Legendre, P., and L. Legendre. 2012. Numerical Ecology. Elsevier, Amsterdam.

Leibold, M. A., E. P. Economo, and P. Peres-Neto. 2010. Metacommunity phylogenetics: Separating the roles of environmental filters and historical biogeography. Ecology Letters 13:1290–1299.

Maestri, R., N. S. Upham, and B. D. Patterson. 2019. Tracing the diversification history of a Neogene rodent invasion into South America. Ecography 42:683–695.

Morrone, J. J. 2014. Biogeographical regionalisation of the Neotropical region. Zootaxa 3782:1–110.

Morrone, J. J. 2015. Biogeographical regionalisation of the world: A reappraisal. Australian Systematic Botany 28:81–90.

Musser, G., and M. Carleton. 2005. Superfamily Muroidea. *in* D. E. Wilson and D. M. Reeder, eds. Mammal species of the world: a taxonomic and geographic reference (3rd ed.). Smithsonian Institution, Washington.

Olivero, J., A. L. Márquez, and R. Real. 2013. Integrating fuzzy logic and statistics to improve the reliable delimitation of biogeographic regions and transition zones. Systematic Biology 62:1–21.

Pielou, E. C. 1984. The Interpretation of Ecological Data. John Wiley, New York.

Pillar, V. D., and L. D. S. Duarte. 2010. A framework for metacommunity analysis of phylogenetic structure. Ecology Letters 13:587–596.

Qiu, Z. D., and C. K. Li. 2003. Rodents from the Chinese Neogene: Biogeographic relationships with Europe and North America. Bulletin of the American Museum of Natural History 586–602.

R Core Team. 2018. R: a language and environment for statistical computing. R Foundation for Statistical Computing, Vienna, Austria.

Ree, R. H., and S. Smith. 2008. Maximum likelihood inference of geographic range evolution by dispersal, local extinction, and cladogenesis. Systematic Biology 57:4–14.

Revell, L. J. 2012. phytools: An R package for phylogenetic comparative biology (and other things). Methods in Ecology and Evolution 3:217–223.

Schenk, J. J., K. C. Rowe, and S. J. Steppan. 2013. Ecological opportunity and incumbency in the diversification of repeated continental colonizations by muroid rodents. Systematic Biology 62:837–864.

Steppan, S., R. Adkins, and J. Anderson. 2004. Phylogeny and divergence-date estimates of rapid radiations in muroid rodents based on multiple nuclear genes. Systematic Biology 53:533–553.

Steppan, S. J., and J. J. Schenk. 2017. Muroid rodent phylogenetics: 900-Species tree reveals increasing diversification rates. PLoS ONE 12:e0183070.

Vellend, M. 2001. Do commonly used indices of B-diversity measure species turnover? Journal of Vegetation Science 12:545–552.

Vilhena, D. A., and A. Antonelli. 2015. A network approach for identifying and delimiting biogeographical regions. Nature Communications 6:1–9.

Wallace, A. R. 1876. The geographical distribution of animals. Harper & Brothers, New York.

Wilson, D. E., and D. M. Reeder. 2005. Mammal Species of the World. A Taxonomic and Geographic Reference (3rd ed). Society 61:2142.

